# QTLIT: A user-friendly Shiny app for Quantitative Trait Loci (QTL) analysis

**DOI:** 10.64898/2025.12.21.694342

**Authors:** Tapan Kumar Mondal, Abhijeet Roy, U B Angadi, Abhishek Mazumder

## Abstract

Identification of Quantitative Trait Loci from DNA sequence data is a multistep process, which is difficult for the plant breeders. To make the QTL identification breeders friendly, we developed QTLIT, a tool comprised of popular R packages and open sources software under a single umbrella for QTL analysis that enables intuitive exploration, multidimensional analyses, and visualization of results generated from any kind of DNA sequence data that deals with Single Nucleotide Polymorphism (SNP) variation. This tool requires raw or processed FASTQ files and proceed it for SNP identification to further perform QTL analysis on genomic regions of interest along with phenotyping data of various pedigree-based mapping populations. QTLIT generates a full range of plots such as VCF quality, linkage map, LOD score etc. It has broad range of applications and options for different parameters which the user can adjust as per their requirements. Moreover, it has two separate interfaces where one can utilize this tool to generate alignment files and/or variant calling for other downstream analysis. QTLIT is a fast and intuitive QTL analysis tool suitable for a wide range of users due to its architecture and its user-friendly graphical interface. QTLIT source code is available at https://github.com/TKM-Lab/QTLIT and the user interface version is available on http://14.139.229.202:8080/QTLIT.

## Introduction

Next-generation sequencing (NGS) technology has enhanced biological research by allowing rapid generation of huge genomic datasets. The genomic dataset comprises base changes, insertions and deletions of nucleotides, large deletions of exons or whole genes and rearrangements such as inversions and translocations. Analysis of this data is essential part to identify and annotate these changes, mapping genomic location to important traits/phenotypes. In principle, Quantitative Trait Locus (QTL) analysis is a statistical association study between phenotype and genotype to know inheritance pattern of quantitative trait and to identify markers applied in breeding using Marker Assisted Selection (MAS). The term Quantitative Trait Locus (QTL) was coined by Gelderman (1919) as a region of the genome harboring polygenes that is associated with an effect on a quantitative trait, controlled by several genes within the specific region. Precise analysis of this process is conventionally done by stepwise analysis using various open source and proprietary softwares. An essential part of the analysis pipeline involves short read alignment and variant calling, the process of identifying genetic variations such as single nucleotide polymorphisms (SNPs) and insertions/deletions (InDels) from raw sequencing reads. While tools like the BWA-MEM (Li and Durbin 2009), Bowtie (Langmead et al. 2009) are popularly used for alignment whereas Genome Analysis Toolkit (GATK) (McKenna et al. 2010), VarScan (Koboldt et al. 2012), Samtools/mpileup (Li 2011) and FreeBayes (Garrison and Marth 2012) for variant calling are in routine use but their command-line interfaces and complex parameter settings often present a barrier for many wet lab researchers. Along with, these tools are individually dedicated to either short reads alignment or variant calling which further hinders the researchers having a streamlined process.

Apart from variant calling, researchers often seek to understand the functional implications of these genetic variations, particularly in the context of phenotypic traits. Most of the phenotypic traits of economic importance are often complex, influenced by both genetics (many genes with varying effects or few genes with substantial effects) and environment. Identifying the genetic regions associated with these traits is a key step in understanding their inheritance for crop improvement/breeding. This involves confirming and refining the location of the markers, usually Single Nucleotide Polymorphisms (SNPs) and/or insertion/deletion (InDel) markers linked with the phenotypic traits in specific chromosomes, known as Quantitative Trait Loci or QTLs, followed by Marker-Assisted selection (MAS). This enabled breeders to select or pyramid favourable alleles for traits like yield, stress tolerance, disease resistance etc. through MAS with efficiency and precision in conventional plant breeding. Moreover, a successful MAS requires accurately mapped QTLs. Precision in phenotyping is important in QTL mapping studies as a few traits are highly influenced by the environmental factors. For precise chromosomal location(s) to obtain from the mapping study in order to identify putative novel candidate genes within specific marker intervals, stable and consistent phenotypic data is essential.

The statistical methods for identification of QTL have been advanced but the simplicity, robustness and efficiency of Composite Interval Mapping (CIM) make it still useful today even though Inclusive Composite Interval Mapping (ICIM) and Genome-wide Composite Interval Mapping (GCIM) have been developed as newer methods. These methods work effectively with marker maps that have limited density and traits controlled by multiple major QTLs because the additional complexity of ICIM or GCIM does not lead to better results. The model used in CIM is simpler than ICIM because it has fewer parameters which decreases the chance of overfitting. The QTL mapping community relies on CIM because it provides a standard method for research that enables comparison with existing published studies from the past several decades. CIM continues to serve as a reliable and acceptable QTL analysis method because it provides efficient results for many genetic research projects even though ICIM and GCIM offer better performance with dense SNP data and complex genetic systems.

Numerous tools are available for linkage mapping and QTL analysis, that are both graphical user interface (GUI) and command-line interface (CLI) based. Among user friendly tools for linkage mapping, JoinMap (https://www.kyazma.nl/index.php/JoinMap/) combined with MapQTL (https://www.kyazma.nl/index.php/MapQTL) is the most popular tools whereas QTL cartographer (Wang et al. 2012) and ICIMapping (Meng et al. 2015) are also a choice for QTL analysis. Even though these tools are user-friendly but they have their own drawbacks, for instance, JoinMap/MapQTL are Windows based and commercial, difficult in handling large data sets and representation of large tables and graphs. Whereas QTL Cartographer is outdated and ICIMapping tool, although free but restricted to Windows. Apart from commercial tools, researchers have developed few tools under open source for analyses like QTL mapping, population dependent linkage analysis and genome-wide association studies including R/qtl (Broman et al. 2003), rTASSEL (Monier et al. 2022), PLINK (Purcell et al. 2007) and GAPIT (Lipka et al. 2012). Further, difficulties in installation due to dependency on other packages, and complex parameter settings often present a barrier for many wet lab researchers globally, especially for researchers having low levels of computational skill, and its command-line interface which can be an uphill task for the beginners. These tools need high performance computing with large storage and memory management system to utilize high throughput data. Moreover, many independent stand-alone tools as a pipeline for variant calling, linkage mapping, and QTL analysis are available. However, a major obstacle is the absence of unification across these pipeline. The conventional method often involves manual data transfer and formatting between many software programs. This not only raises the risk of error incorporation but also leads to retardation of the research program. Moreover, these analysis need high computational power and specialized computational skills. Many of these programs require scripting expertise to use from the command line interface and does not have interactive graphical v isualization that also depend on a specific operating system. Furthermore, the steep learning curve related with becoming proficient in these programs might deter researchers with limited knowledge in bioinformatics.

To address these challenges, we developed a web-based R shiny server tool QTLIT, and deployed on dedicated high performance computing system that performs comprehensive QTL analyses with visual interactive and graphical user interface on different user requirements/inputs. Therefore, QTLIT will enable non-expert bioinformatics researchers to directly perform these analyses in a rapid and user-friendly manner.

## Materials and methods

### R packages

The QTLIT web tool has been developed with R programming language using various R-package libraries using shiny server with web interface. The “shiny” package was developed (Chang et al. 2025) in order to enable developers to combine R codes with computer languages like HTML and JavaScript and thereby make it more accessible via web browsers through internet. Shiny server supports the extensive library of R packages for user-friendly GUI interface. Currently, implemented in computing power system with 128 GB RAM and 32 cores for processing in LINUX environment for the tool to perform the tasks.

This involves, Variant Calling and subsequent QTL analysis through various R packages such as : rtracklayer (Lawrence et al. 2009) for import/export of fasta files, ShortRead (Morgan et al. 2009) for quality check of raw FASTQ files, gmapR (Wu et al. 2016) for reference genome indexing and short reads alignment, Rsamtools (Morgan et al. 2024) for indexing binary alignment (BAM) file, vcfR (Knaus and Grünwald 2017) for VCF plotting, Biostrings (Pagès et al. 2024) for manipulation of large sequences, stringr for pattern matching, rvest for manipulation of HTML file, ggplot2 for enhanced plotting, rTASSEL (Monier et al. 2022) for VCF filteration, ASMap (Taylor and Butler 2017) for linkage map construction, LinkageMapView (Ouellette et al. 2018) for linkage map generation, R/qtl (Broman et al. 2003) for QTL analysis, gridExtra (Auguie 2017) for placing multiple plots in a grid, parallel for parallelization of the process, qtlhot (Neto and Yandell 2018) for phenotype input and ggrepel (Slowikowski 2024) for labelling of the significant markers. All the packages are tested on RStudio (v 2023.12.1-402). The version of the packages and other dependencies are available in supplementary file, a custom R script was also incorporated for the conversion of VCF file to R/qtl cross object (https://github.com/RimGubaev/v_to_qtl/blob/master/vcf_to_qtl_converter.R) in the shiny app scripting. Apart from the R packages, bcftools (v 1.13) (Danecek et al. 2021) was utilized for the variant calling utilizing system command in the R environment.

### Workflow and methodology

The complete implementation of QTL, as show in Fig. 1, encompasses two major steps (a) Variant analysis and (b) Linkage map and QTL analysis. The variant calling tab proceeds with read alignment, filtrations and variant call, whereas, the QTL analysis tab includes linkage map construction and QTL mapping.

**Figure 1.**
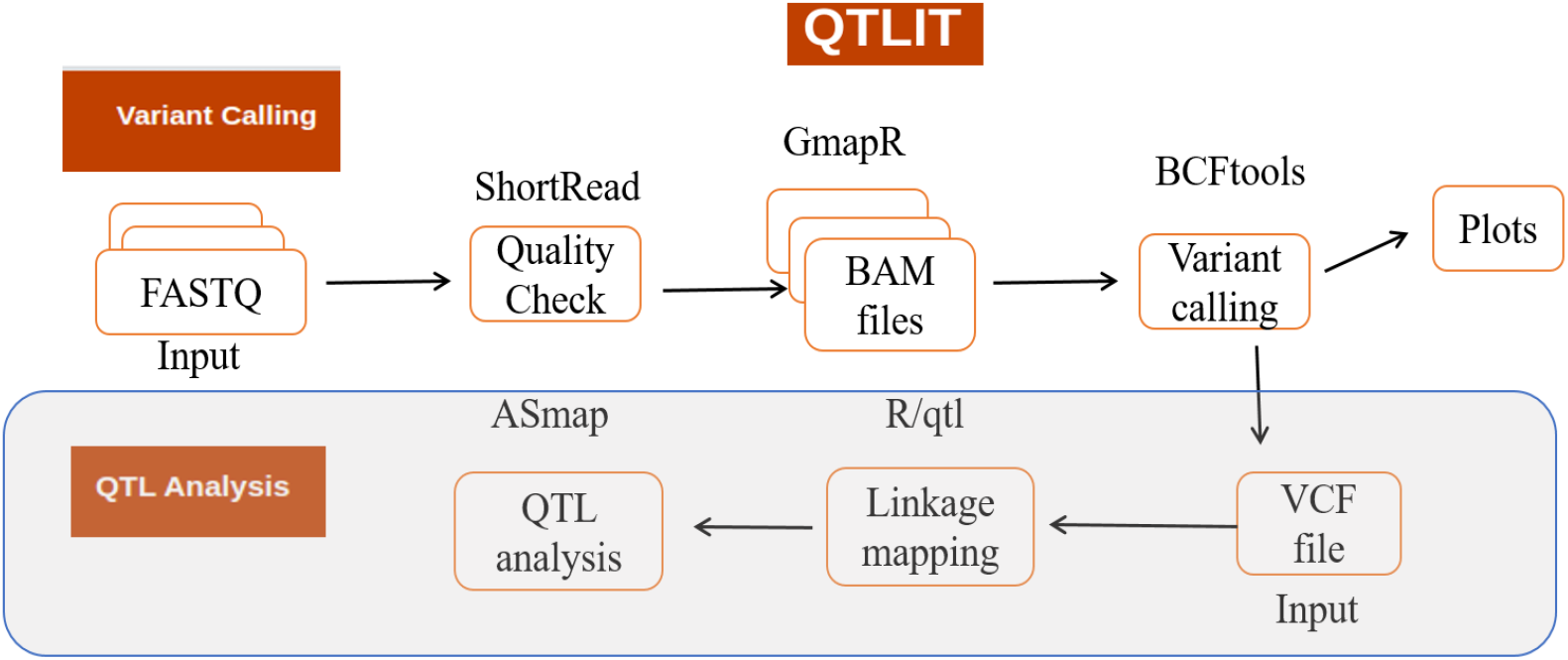
Pipeline of the QTLIT tool from variant calling to QTL analysis and the packages used in developing this tool.

### Program overview

QTL Identification Tool (QTLIT) is multi-functional GUI web tool, which generates QC report, binary alignment file (BAM), variant plots and variant calling file (VCF). The VCF file is further utilized for the linkage mapping and QTL identification with user defined parameter s. QTLIT pipeline follows two individual workflow dedicated to (a) variant calling analysis which is divided into two sub-sections: (1) Short read alignment and generation of the BAM files, (2) Variant calling and generating the VCF plots and result file for download, (b) Linkage mapping and QTL identification with a user-friendly interface having separate panes for parameters and tabs for result display/downloads. Users can upload VCF and phenotype files, and set parameters. Analysis execute at various steps of the pipeline, where the results of every step will be used for subsequent analysis/step with a progress log and generated plots.

### (a) Variant analysis

In variant calling process, first step is to generate BAM files for which reference genome fasta fil e either in zipped or unzipped format is required which will be auto indexed using the gmapR’s *GmapGenome* function with the default settings of base size 12 and K-mer size of 15. Further, user can upload multiple paired-end raw or processed fastq files of genotypes, the uploaded file can be proceeded for alignment with options by skipping the Quality Checking or assess the quality of the short reads before proceeding for the alignment. In Alignment, the paired end short reads are aligned to the indexed reference genome using gmapR, *gsnap*function with default options. Then, variant calling is to be performed using BCFTools *mpileup* and *call* function with mapping and base quality filter of 20 and filtration quality >30 for PASSED-flagged biallelic SNPs through BCFtools *view* function. Each of the individual variant files are then automatically merged for downstream analysis.

### (b) Linkage map and QTL analysis

For linkage map construction and subsequent QTL analysis, the workflow follows a standard QTL mapping procedure, which comprises several filtering and pre-processing. Alongwith input files, the user can specify the parental alleles and the type of phenotype to be analyzed. A dynamic user interface element allows the user to select column containing the phenotype data from the uploaded CSV file. Several filtering steps are implemented as default to ensure better data quality while proceeding for next steps. These include filtering markers based on minimum site count i.e. the minimum number of count which is known as allele frequency, is set to minimum of 5% (0.05) and maximum of 100% (1.0) but the user can change based on the needs alongwithheterozygosity with default minimum value of none and maximum of 10% (0.1). An option is also provided to remove minor SNP states based on less common allele frequency. Markers which are exhibiting segregation distortion with p-value threshold of 0.05 or beyond, a user-defined threshold will be dropped in the filtering process and plot generation.

While in linkage mapping, the user can select from different mapping functions (Haldane and Kosambi) for the linkage map and two more i.e., Carter-Falconer and Morgan mapping function, to convert recombination frequencies to genetic distances. Also, p-value for linkage clustering can be set based on the population size and the number of markers, which is currently set to 1.0 that the user can change in the interface.

The users have an option for QTL analysis using either a single-locus scan (scanone) or composite interval mapping (CIM) approach alongwith selecting the options for other parameters defined in the QTLIT graphical user interface (GUI). Moreover, based on the analysis models, options (scanone or CIM) can also be set by the user for calculating the test statistics (EM, imputation, Haley-Knott regression or efficient Haley-Knott regression). The output files of this analysis will be a summary statistics of the phenotypic data displayed as the text output, a set of plots summarizing the distribution of the selected phenotype along with a download option for the linkage map plot generated in pdf format in the GUI and a LOD plot generated either from scanone or CIM results grouped with genotype and marker distribution plots from the QTL analysis.

## Results

### Data inputs and analysis

The developed web tool is customized with three levels of inputs and analysis, (1) User can initiate with raw data of sequences such as reference genome and multiple genotypes in FASTA or compressed fasta.gz format for BAM file generation (2) Aligned BAM files for variant calling and VCF file generation (3) VCF files with phenotype file for linkage mapping and QTL analysis.

### Graphical User interface

The QTLIT tool has two tabs with each tab having side menu and dedicated analysis button. For variant calling tab, side menu is divided into uploading the files viz. (1). Reference Genome file in FASTA format (2) Paired-end genotype FASTQ file or (3) BAM file either generated from the analysis or already user prepared BAM files with analysis buttons for BAM file generation and Variant Calling. In QTL analysis tab, the side pane consists of uploading option for two files viz. (1). VCF file generated from the tool or user prepared VCF and (2) Phenotype file in CSV format, and a single Run Analysis button. The visualization of workflow progress can be seen in the result display. After completion of the task, the results can be downloaded in the individual tab sections with sub-tabs in variant calling analysis for QC report, BAM file, VCF file and VCF plots and from the four sub-tabs in Linkage mapping and QTL analysis which includes a data summary, phenotype data summary, pdf file of linkage map and QTL analysis results (Fig. 2). At each level of analysis, user friendly parameter setting has been provided with default values.

**Figure 2.**
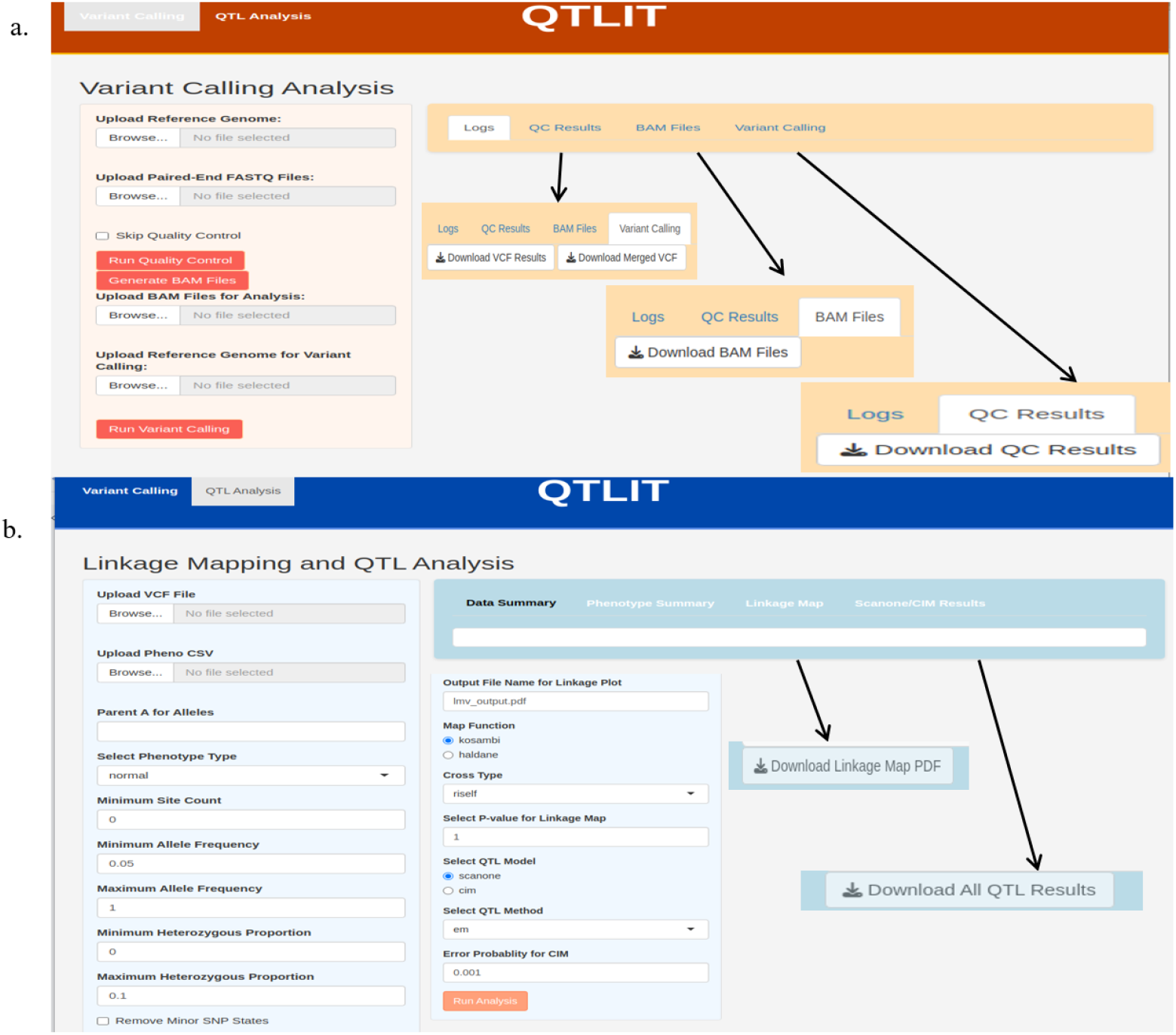
User interface of QTLIT tool (a) Variant calling analysis with side bar for uploading reference genome, bam files and analysis running button and each of the analysis having a dedicated tab for downloading results, (b) QTL analysis console with side bar included in the inlay depicting different selection parameters.

### Variant calling

An intuitive workflow for variant calling is integrated in QTLIT, enabling users to further process raw or processed sequencing data from FASTQ files to generate a variant calling file in VCF format with different breakpoints. Immediately after uploading the uncompressed reference genome for indexing of the FASTQ files, if the user initially desires to check the quality and statistics of the FASTQ files that provide a result folder in zipped format consisting of image files, pdf file and html report of the quality assessment including base calls statistics, per cycle base calls, read counts, read occurrence and read quality. As soon as user clicks on the “Generate BAM files”, auto indexing of reference genome initiates with the default parameters and further used for alignment, generating the BAM files which users can download from the BAM files sub-tab. The users can utilize this module for generating the BAM files for other analysis including transcriptomic analysis, if the FASTQ files are RNA-seq data instead of DNA-seq. Also, an option is provided to upload the BAM files for variant calling, if users already have BAM files with them generated from different softwares or else user can go to the next step. The tab “Upload Reference Genome for Variant Calling” is explicitly provided and the reason being, in the next step of variant calling the same reference genome in uncompressed format needs to be re-uploaded to be indexed by BCFtools. Therefore, instead of automating the direct variant calling, this break point was provided for having an option to upload already generated BAM files from some other source and to directly proceed with the variant calling after uploading the reference genome file in an unzipped form. The main panel displays a log tab of the workflow progress as well as zipped files can be retrieved from respective tab for further analysis, in which “Variant Calling” tab consists of the plot folder and a merged VCF file in compressed form, completing the end-to-end variant calling pipeline. Currently, the plots of variant quality and Depth of coverage (DP) distribution is getting generated, which depicts a visual representation of the counts quality score and read depth of coverage, respectively, (Fig. 3) that users can download from the console apart from the VCF file in compressed format for linkage mapping and QTL analysis.

**Figure 3.**
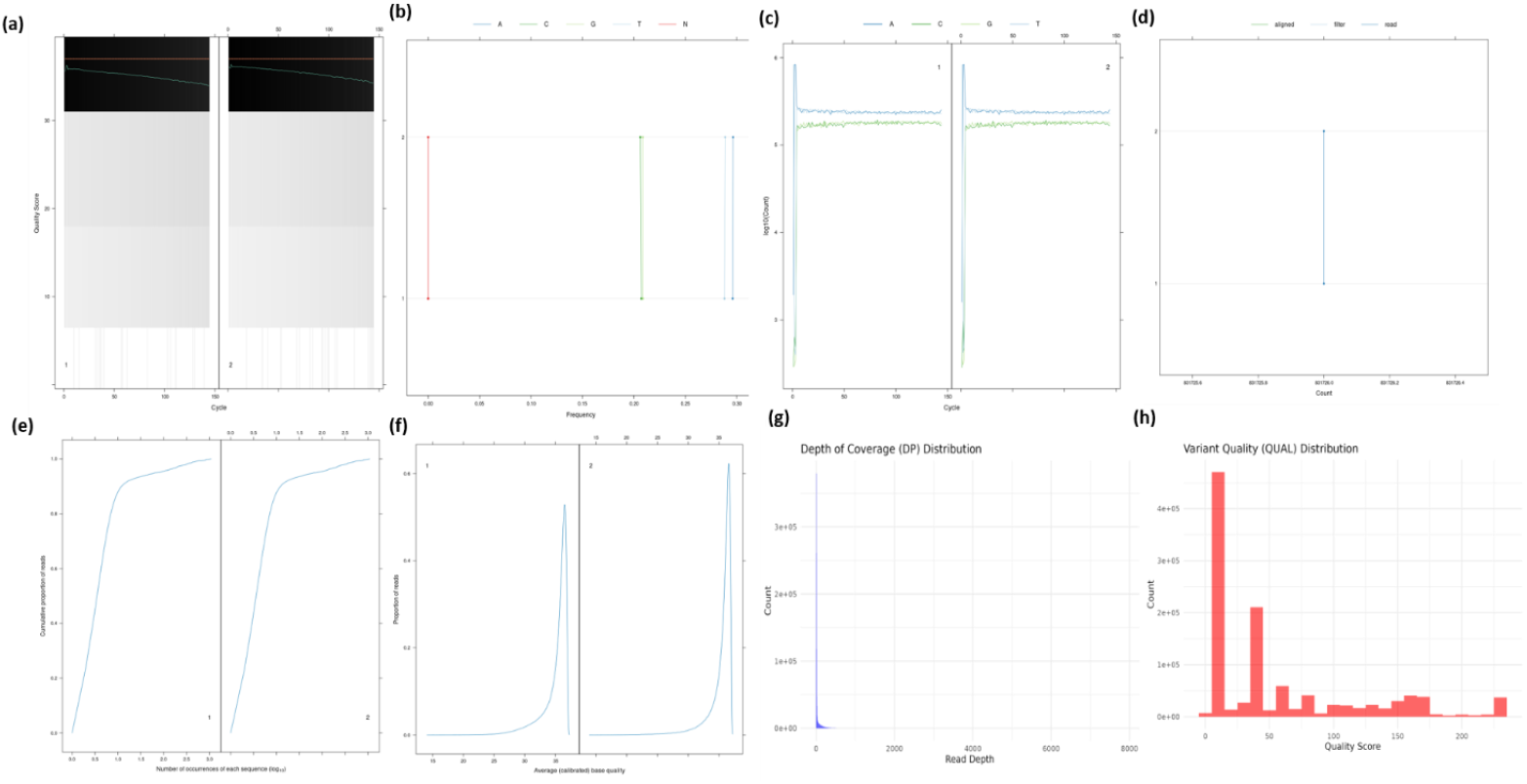
VCF calling pipeline results (a) quality score per cycle, (b) frequency of bases, (c) base counts per cycle, (d) average number of reads, (e) cumulative proportion of reads and the number of sequences, (f) proportion of reads with average base quality, (g) depth of coverage in the VCF file, (h) count wise variant quality score. (VCF: Variant Calling Format)

### Linkage mapping and QTL analysis

This section is the second and final part of the QTLIT tool including various setting and parameter options for the analysis. Starting with the uploading of unzipped VCF file generated in the previous step and the phenotype information in the CSV format, in which users have to specify the type of phenotypic data and selecting the desired phenotype column of phenotype data for analysis, if multiple phenotype columns are present. The users also have to provide parent A name for allele coding into ABH (A for homozygous parent A, B for homozygous parent B and H for heterozygous) format for R/qtl cross object. The sidebar allows users to configure the parameters, including filtering criteria for genotypic data (minimum site count, allele frequencies, and heterozygous proportions), segregation distortion thresholds, and parameters for linkage mapping (map function, cross type). The main panel displays the results in different tabs which users can visualize after toggling, including a linkage map plot generated in a pdf format using clustering algorithm MSTmap (Wu et al. 2008), and the map can be downloaded from the respective section. A phenotype data summary, Phenotype summary plots (histograms, boxplots, QQ plots), are also getting generated in the respective tabs (Fig. 4). Whereas QTL mapping results (either scanone or CIM) which include markers vs. individuals and vice-versa, generate genotype comparison and a LOD score plot as well (Fig. 5).

**Figure 4.**
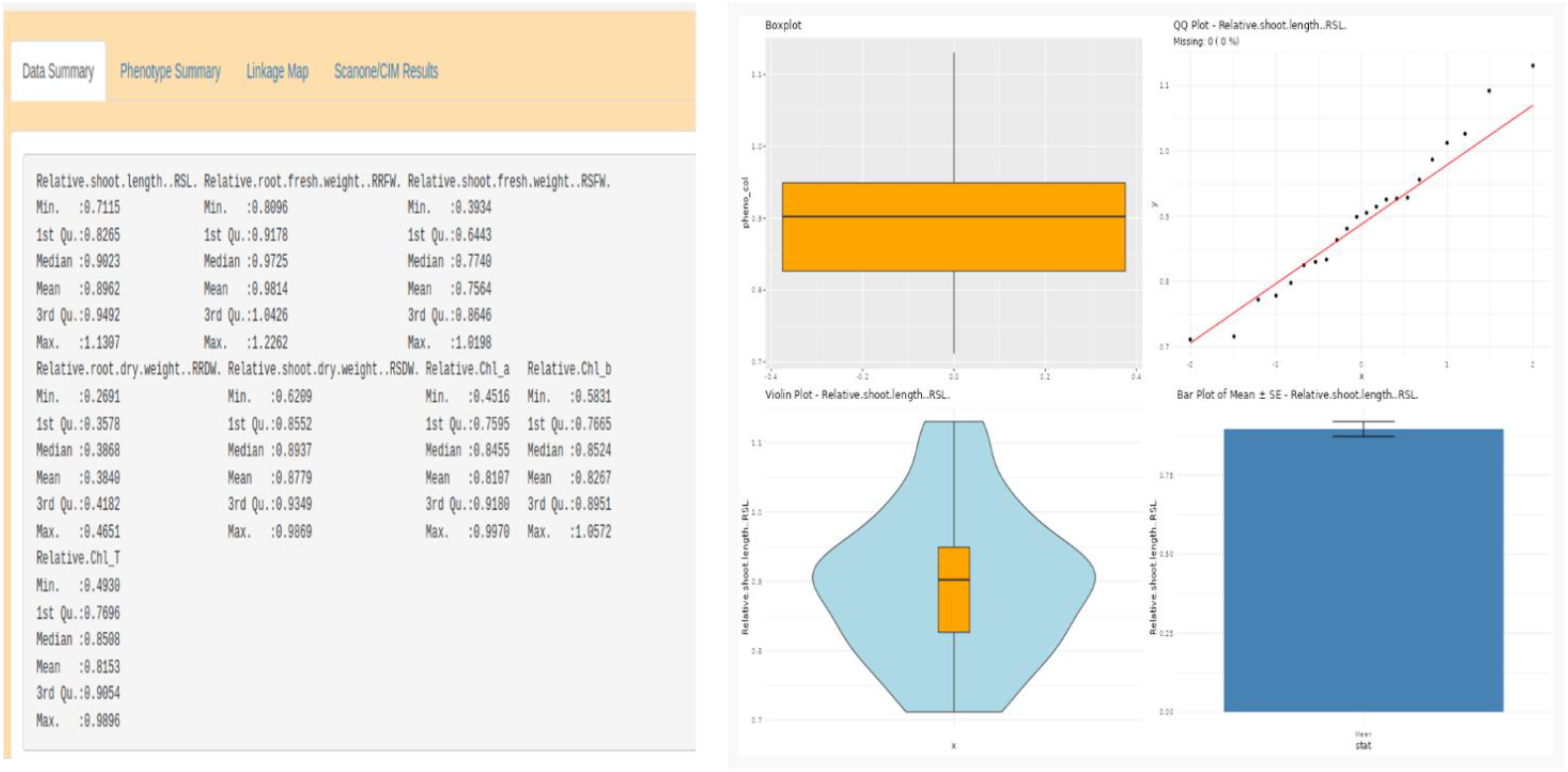
Result outputs of the phenotype data provided by the users, (a) data summary of all the multiple phenotypes information provided by the user, (b) box plot, Q-Q plot, violin plot and mean with SD of the selected phenotype. (SD: Standard deviation, Q-Q: Quantile-Quantile)

**Figure 5.**
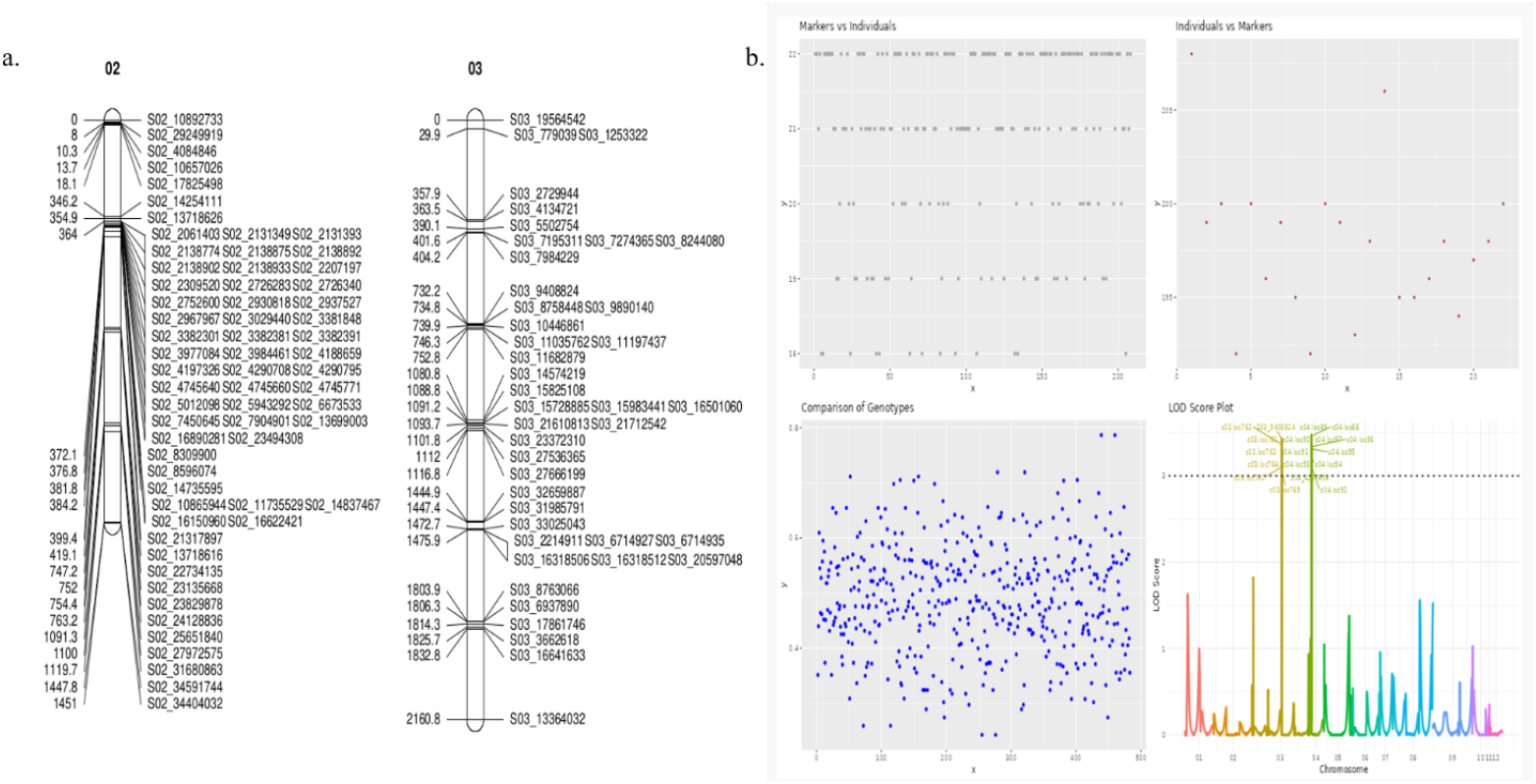
Result summary of the QTL analysis, (a) a snapshot of the linkage map generated in the analysis, (b) various plots of QTL analysis, genotype comparison and LOD plot with significant markers being labelled.

## Discussion

The QTLIT provides a comprehensive, user-friendly interface, integrated with key genomic analysis workflows: Short reads alignment, variant calling, linkage mapping, and finally QTL analysis into a single window solution, by addressing a notable gap in marker-trait association studies. As most of these tools available are individually dedicated to variant calling, linkage mapping and/or QTL analysis. A unified framework that connects the workflow remains a challenge due to unification of inputs from one tool to another. This leads to hindered efficiency, end-to-end genomic investigations and particularly in research areas where computational skills of the user is low. Moreover, the requirement of computational skill for command line based tools is a significant hurdle for non-bioinformatics users to utilize diverse tools having complex parameter settings and command line interface. Variant calling and QTL analysis require command line skills and computational power. The developed web tool addresses all the aforementioned issues.

The tool incorporates packages which are highly efficient and regularly updated where the function *gsnap* in gmapR is known for its accuracy and efficiency in aligning the short reads to large genomes (Wu et al. 2016). As the importance of accuracy in the alignment for variant calling is very crucial as highlighted in the study by Li & Durbin, 2009, where it revealed the impact of alignment algorithms on downstream variant calling accuracy. Further, we are more focused on plant breeding and the BCFtools (Li 2011), variant calling is better for non-human and non-model subjects as it generates lower number of false positives (Lefouili and Nam 2022). For, QTL analysis, the open source package R/qtl (Broman et al. 2003) can be employed but the steep learning curve of R programming makes it not to be chosen over other GUI based QTL analysis tools, which has been addressed here by integrating parameter setting meticulously used in the QTL analysis. The tool supports both single-QTL scan (*scanone*) and composite interval mapping (*cim*) models and flexibility in QTL detection. The recombination frequency calculation for linkage mapping using MSTmap algorithm (Wu et al. 2008) is integrated with ASMap tool (Taylor and Butler 2017) which uses minimum spanning tree algorithm (Cheriton and Tarjan 1976) for clustering and this is much faster and efficient for the large datasets which is deficient in *est*.*rf* function of R/qtl. This feature also has been addressed here. Apart from ASMap and R/qtl, only R package available for linkage mapping is OneMap (Margarido et al. 2007) but it is restricted to outbred and inbred populations. Moreover, an integrated R package LinkageMapView (Ouellette et al. 2018) to visualize the linkage map in high resolution is aesthetically appealing. This will avoid the dependency on other visualization tool like windows based tool MapChart (Voorrips 2002) . Therefore, integrating all these packages, QTLIT simplifies the complex workflow associated with identifying genomic regions or the SNP markers influencing the phenotypic traits. Moreover, the overall statistics of the phenotype data can also be assessed in the single window.

In summary, QTLIT provides a valuable platform for variant calling, linkage mapping and QTL analysis, addressing critical aspects of data preprocessing, analysis, and visualization. Its user-friendly interface makes it a valuable resource for researchers in plant breeding as well as in diverse fields.

## Data availability

https://github.com/TKM-Lab/QTLIT

## Acknowledgement

The authors are grateful to the Director, National Institute for Plant Biotechnology (NIPB), New Delhi for providing all the necessary facilities to execute the research work.

## Author’s contributions

AR wrote the major portion of the manuscript including the pipeline and prepared the initial draft, UBA improved the pipeline and manuscript, AM assisted in the QTL visualization and inputs, TKM conceptualized the idea, improved the manuscript, and helped in the fund acquisition.

